# RACK1 is required for axon guidance and local translation at growth cone point contacts

**DOI:** 10.1101/816017

**Authors:** Leah Kershner, Taylor Bumbledare, Paige Cassidy, Samantha Bailey, Kristy Welshhans

**Affiliations:** Department of Biological Sciences, Kent State University, Kent, Ohio, USA; School of Biomedical Sciences, Kent State University, Kent, Ohio, USA; Brain Health Research Institute, Kent State University, Kent, Ohio, USA

## Abstract

Local translation regulates the formation of appropriate connectivity in the developing nervous system. However, the localization and molecular mechanisms underlying this translation within growth cones is not well understood. Receptor for activated C kinase 1 (RACK1) is a multi-functional ribosomal scaffolding protein that interacts with β-actin mRNA. We recently showed that RACK1 localizes to and regulates the formation of point contacts, which are adhesion sites that control growth cone motility. This suggests that local translation occurs at these adhesion sites that are important for axonal pathfinding, but this has not been investigated. Here, we show that RACK1 is required for BDNF-induced local translation of β-actin mRNA in growth cones. Furthermore, the ribosomal binding function of RACK1 regulates point contact formation, and axon growth and guidance. We also find that local translation of β-actin occurs at point contacts. Taken together, we show that adhesions are a targeted site of local translation within growth cones, and RACK1 is critical to the formation of point contacts and appropriate neural development. These data provide further insight into how and where local translation is regulated, and thereby leads to appropriate connectivity formation in the developing nervous system.

## Introduction

The establishment of neural connectivity requires the precise regulation of axon guidance during development. At the tips of developing axons are highly dynamic and motile mechanosensory structures known as growth cones. Growth cones contain filopodia, finger-like protrusions, in their leading edge, and lamellipodia, sheet-like veils, between filopodia. Both filopodia and lamellipodia are composed primarily of filamentous actin (F-actin) and contain receptors that bind substrate-bound and diffusible guidance cues in the extracellular environment. Guidance cue binding leads to activation of intracellular signaling cascades, which ultimately results in changes in cytoskeletal dynamics and growth cone steering toward their proper targets (1).

Receptor for activated C kinase 1 (RACK1) is a seven WD40 domain ribosomal scaffolding protein that is a member of many signaling pathways (2, 3). It is crucial for overall development as mice lacking RACK1 are gastrulation lethal (4). Recent research has shown the importance of RACK1 specifically in neural development. RACK1 regulates neural tube closure in *Xenopus* (5), axon guidance in *Caenorhabditis elegans* (6), and initial neurite outgrowth in PC12 cells (7). Furthermore, our recent study shows that RACK1 regulates axon outgrowth of primary mammalian cortical neurons (8). Together these studies suggest an important role for RACK1 in neural development; however, the mechanisms underlying this role are unknown. Here, we find that RACK1 regulates axon outgrowth and guidance. Because binding between RACK1 and ribosomes is critical for neural development, we hypothesized that RACK1’s main role in this process is the regulation of local translation.

Local translation is the process by which proteins are translated where they are spatially needed rather than being translated in the soma and then transported to subcellular regions. Locally translated mRNAs contain binding motifs, typically in their untranslated regions (UTRs), that are recognized by RNA-binding proteins (RBPs). For example, zipcode binding protein 1 (ZBP1) is an RBP that recognizes a binding motif in the 3’UTR of β-actin mRNA. The RBP and its bound mRNA are trafficked to the appropriate subcellular location, and after stimulation with specific signaling cue(s), the mRNA is released so that it can be translated (9). RACK1 is known to interact with ZBP1 (10). Furthermore, local translation of β-actin mRNA in growth cones is required for appropriate axon guidance (11, 12). Here, we show that RACK1 is required for BDNF-induced local translation of β-actin mRNA in growth cones. Therefore, RACK1 is critical for local translation and neural development.

Recently, we discovered that RACK1 localizes at growth cone point contacts and is necessary for their formation (8). Because RACK1 is involved in the local translation of β-actin mRNA and localizes to point contacts, we hypothesized that point contacts may be a site of local translation in growth cones. Point contacts, which are somewhat similar to focal adhesions found in other cell types, are integrin-dependent adhesion sites that are comprised of many proteins such as paxillin, talin, vinculin and β1 integrin (13, 14). They stabilize new growth cone protrusions by linking intracellular actin and the extracellular matrix. The mechanisms through which point contacts regulate neurite growth have been examined (15–21). In brief, point contacts are dynamic structures that continually assemble and disassemble during the process of growth cone extension and steering. Here we provide the first evidence that point contacts are sites of local translation. Previous studies have shown that local translation is compartmentalized within axons and dendrites (22), and experiments in mouse embryonic fibroblasts reveal “hotspots” of mRNA and ribosome localization at focal adhesions (23). Like focal adhesions, point contacts are significant regulators of cell motility and now we show that adhesions are key sites of local translation. Because we show that local translation occurs at point contacts, this has strong implications for adhesion mechanisms in general, and certainly warrants further study into how local translation and cell adhesion act not only independently, but also synergistically to direct motility and development.

## Results

### RACK1 is necessary for axon guidance

Here we investigated if RACK1 is required for axon guidance, which is the pathfinding of growth cones in response to a guidance cue. We used the axon guidance Dunn chamber assay (24) and measured the turning angles of individual axonal growth cones to a gradient of BDNF, which is an attractive cue for this neuronal type. Growth cones from neurons transfected with a non-silencing control shRNA turn towards the BDNF gradient, but growth cones from neurons transfected with an shRNA to knockdown RACK1 do not (Figure 1A). When the turning angle was quantified, transfection with RACK1 shRNA resulted in a significant decrease in turning angle, as compared to non-silencing control (Figure 1B). Therefore, RACK1 is required for growth cone steering towards the attractive cue, BDNF.

**Figure 1:**
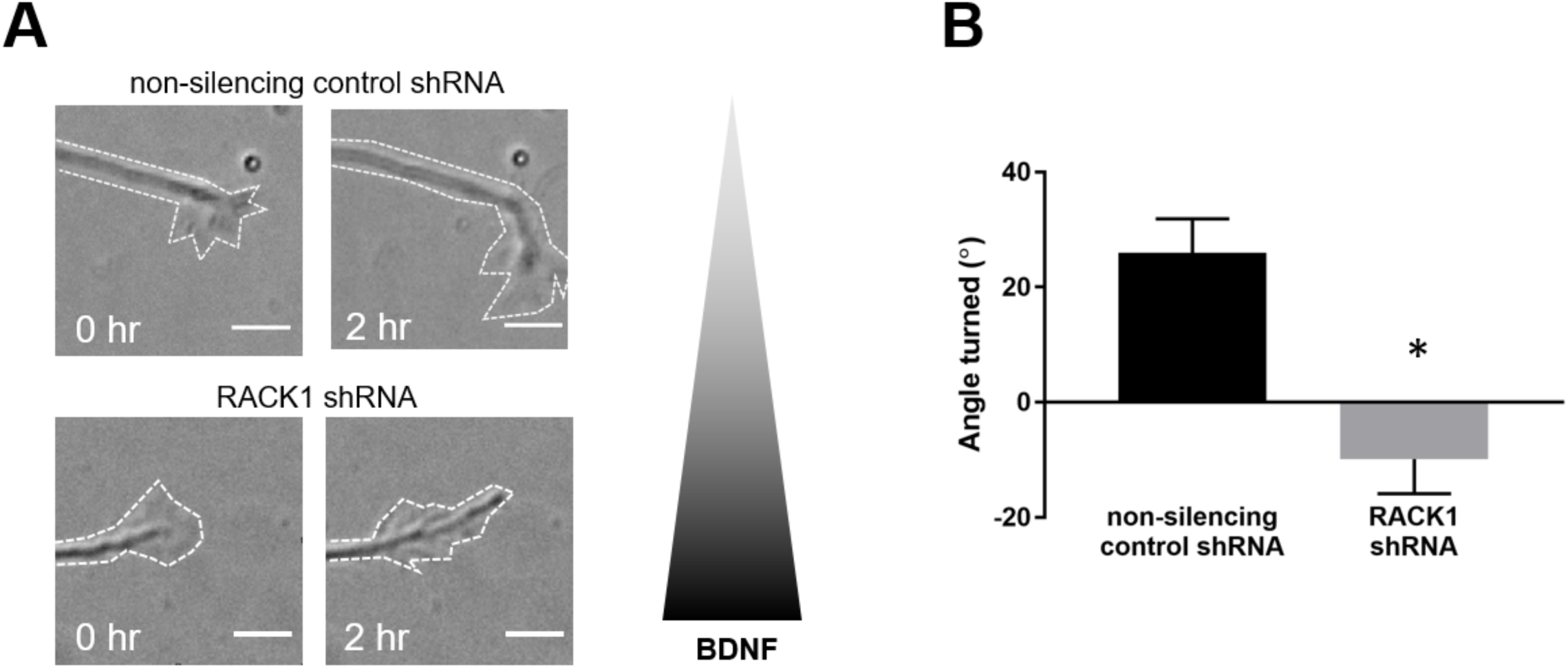
RACK1 is required for growth cone steering towards BDNF. (A and B) Cortical neurons expressing either non-silencing control shRNA or RACK1 shRNA were used in the Dunn chamber turning assay. The outer well of the Dunn chamber contained BDNF, which created a chemoattractive gradient. The triangle to the right of the images indicate the BDNF gradient, such that highest concentrations of BDNF occur at the bottom of each image. Turning angles towards the gradient were measured for the two groups. Scale bars, 5 µm. \**p* ≤ 0.05, Student’s t-test. Non-silencing control shRNA, *n* = 19; RACK1 shRNA, *n* = 11.

### RACK1 is required for BDNF-induced local translation of β-actin in growth cones

Local translation is a common mechanism underlying appropriate axon guidance (reviewed in 9). Because RACK1 is a highly conserved ribosomal scaffolding protein (reviewed in 25) and required for axon guidance (Figure 1), we hypothesized that the underlying mechanism by which RACK1 controls axon guidance might be local translation. Mouse cortical neurons were transfected with either non-silencing control shRNA or a verified RACK1 shRNA (8), and then plated on glass coverslips. At 2 DIV, cells were treated with BDNF, which stimulates local translation of β-actin (11), or vehicle, and then fixed. Quantitative immunocytochemistry was performed to label β-actin protein. Measurements of fluorescence intensity within growth cones of cortical pyramidal neurons transfected with non-silencing control shRNA demonstrated that BDNF results in a significant increase in β-actin. However, shRNA to RACK1 eliminates this increase, suggesting that RACK1 is required for the BDNF-induced local translation of β-actin (Figure 2, A-B).

**Figure 2:**
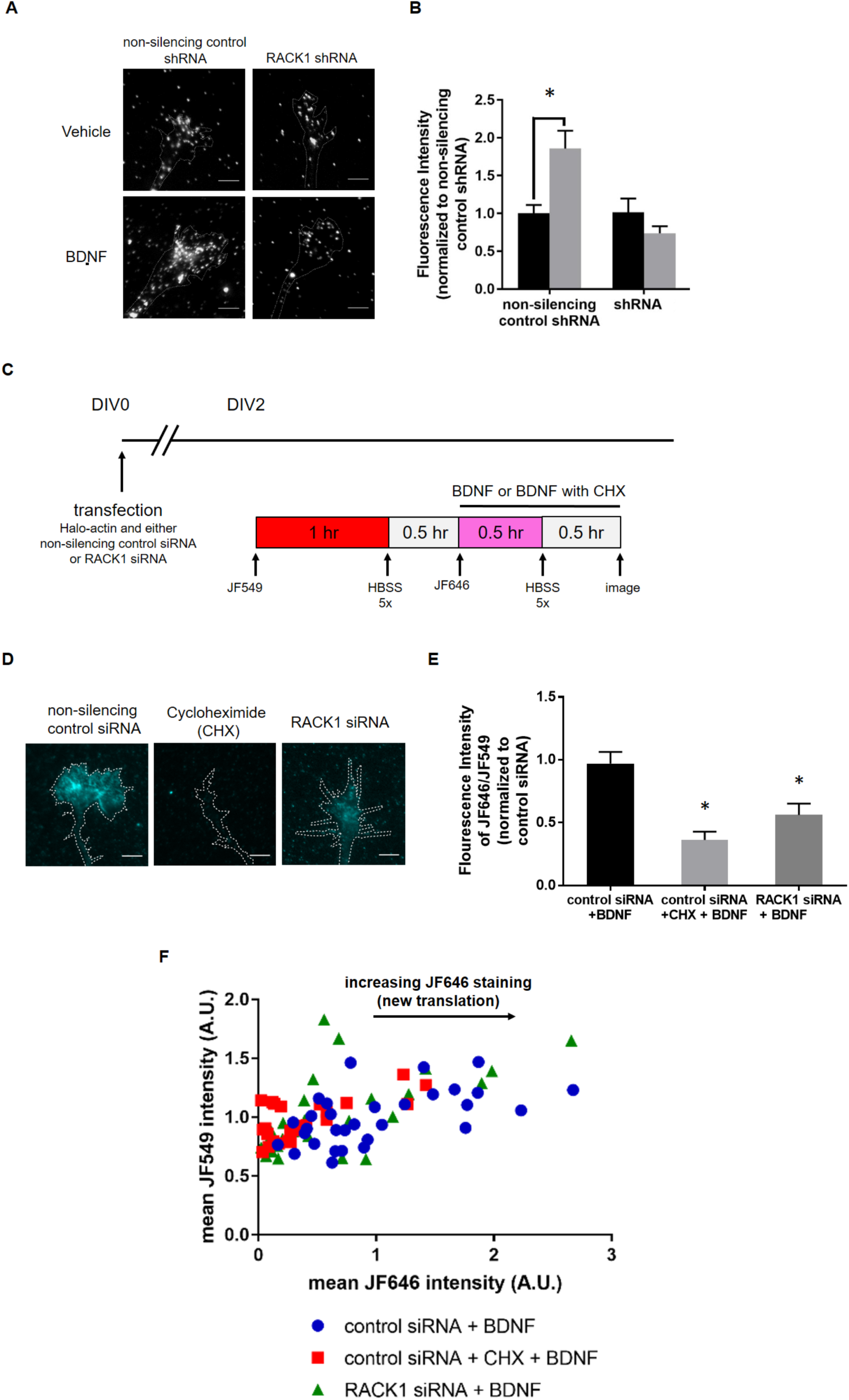
RACK1 is required for BDNF-induced translation of β-actin in growth cones. (A and B) Neurons expressing either non-silencing control shRNA or RACK1 shRNA were exposed to either vehicle or BDNF for 30 minutes. BDNF stimulation results in a significant increase in β-actin protein in neurons expressing non-silencing control shRNA, whereas this increase is lost following knockdown of RACK1 with shRNA. Dotted lines outline axonal growth cones. Note that dotted lines also include the axon for ease of visualization, but axons were not included in the analyses. Scale bars, 5 µm. (B) Quantification of β-actin fluorescent staining in growth cones. All groups were normalized to growth cones expressing non-silencing control shRNA and treated with vehicle. \**p* ≤ 0.05, two-way ANOVA with Dunnett’s multiple comparisons. Non-silencing control shRNA + vehicle, *n* = 60; non-silencing control shRNA + BDNF, *n* = 60; shRNA + vehicle, *n* = 60; shRNA + BDNF, *n* = 60. (C) Schematic of experimental design for D-F. Prior to plating, neurons were transfected with Halo-actin and either nonsilencing control siRNA or RACK1 siRNA. At 2 DIV, neurons were sequentially labeled with JF549 HaloTag ligand prior to BDNF treatment and JF646 HaloTag ligand during BDNF treatment to label preexisting and new translation, respectively. An additional group was transfected with Halo-actin and nonsilencing control siRNA and treated with CHX during BDNF stimulation. (D) Treatment with CHX or knockdown of RACK1 with siRNA significantly reduce translation of β-actin following BDNF stimulation. In the images, nascent JF646-labeled Halo-actin is shown. Dotted lines outline axonal growth cones. Note that dotted lines also include the axon for ease of visualization, but axons were not included in the analyses. Scale bars, 5 µm. (E) Quantification of nascent translation (JF646) over preexisting translation (JF549) for the three groups. \**p* ≤ 0.05, ANOVA with Dunnett’s multiple comparisons. (F) Scatter plot showing JF549 staining (preexisting translation) on the Y axis and JF646 (new translation) on the X axis. Increasing distance on the X axis indicates increased new translation. Non-silencing control siRNA + BDNF, *n* = 31; non-silencing control siRNA + CHX + BDNF, *n* = 24; RACK1 siRNA + BDNF, *n* = 31.

Next, we used a translation reporter to examine whether RACK1 regulates local translation of β-actin in growth cones of living neurons. The HaloTag β-actin reporter is a construct encoding HaloTag (26), followed by the β-actin coding sequence and the β-actin 3’UTR, and is termed “Halo-actin” herein for simplicity. This construct has been used previously to detect newly translated β-actin protein in dendrites (27). Following construct expression, a wide variety of cell permeable fluorophore ligands can covalently bind to the HaloTag which allows for its visualization in live cells (28). Here, we use the red JF549 ligand and the far-red JF646 ligand sequentially to label Halo-actin in live cells. This sequential labeling enables us to distinguish between new and preexisting Halo-actin protein (Figure 2C). Mouse cortical neurons were transfected with the Halo-actin reporter and either non-silencing control siRNA or a RACK1 siRNA and then plated. At 2 DIV, cells were incubated for 1 hour with 20 nM JF549 HaloTag ligand to label preexisting β-actin protein, and then neurons were washed for 30 minutes to remove excess unbound JF549 ligand. Next, neurons were incubated with 200 nM JF646 HaloTag ligand and BDNF for 30 minutes to label nascent β-actin protein, and then washed with HBSS containing BDNF for 30 minutes to remove excess unbound JF646 ligand. We then used live cell imaging to measure the amount of preexisting and BDNF-stimulated nascent β-actin in growth cones. To ensure that Halo-actin was properly reporting translation, in a separate group we incubated neurons transfected with the Halo-actin reporter and non-silencing control siRNA with a protein synthesis inhibitor, cyclohexamide (CHX). When CHX was added at the same time as BDNF stimulation, there was a significant reduction in the translation of Halo-actin versus vehicle control (Figure 2, D-F; compare “control siRNA + BDNF” to “control siRNA + CHX + BDNF”). Furthermore, knockdown of RACK1 significantly inhibited translation of nascent Halo-actin protein (Figure 2, D-F; compare “control siRNA + BDNF” to “RACK siRNA + BDNF”). Taken together, these two experiments demonstrate that RACK1 is required for BDNF-induced translation of β-actin in growth cones.

### Ribosomal binding function of RACK1 regulates neural development

RACK1 is a multifunctional ribosomal scaffolding molecule that can bind to many proteins concurrently. Specifically, RACK1 regulates translation by recruiting kinases, initiation factors, and mRNAs to the ribosome (25). Here, we investigate whether the ribosomal binding function of RACK1 is required for neural development. We first investigated the association between RACK1 and ribosomes under both basal and stimulated conditions. Because RACK1 is required for BDNF-induced local translation of β-actin in growth cones (Figure 2), we expected that BDNF application may initiate local translation by upregulating ribosome binding to RACK1. Using double label immunocytochemistry, we find that RACK1 colocalizes with the 40S ribosomal subunit in growth cones, but this colocalization does not increase following BDNF stimulation (Figure 3, A-B).

**Figure 3:**
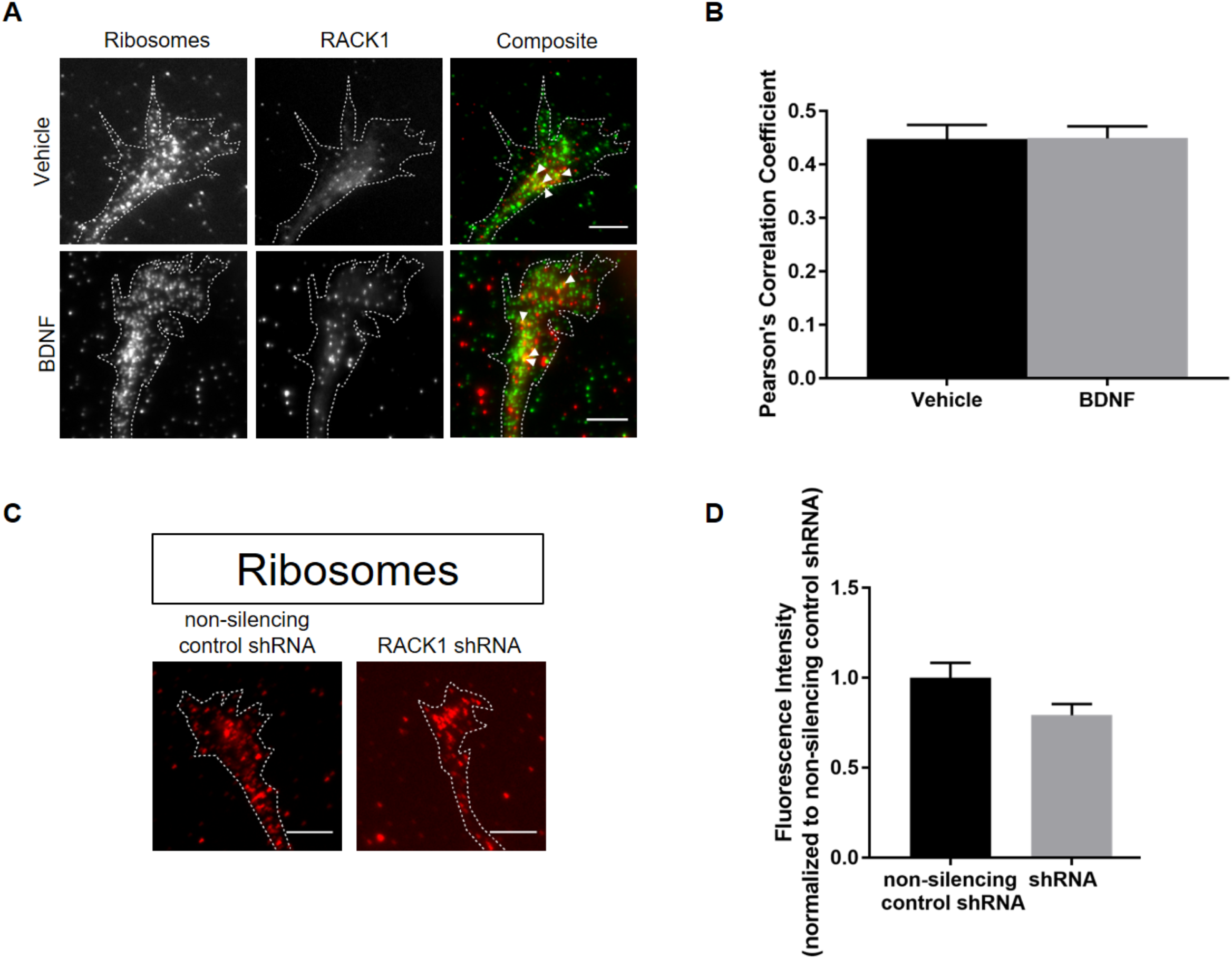
Association between RACK1 and ribosomes is unaffected by BDNF stimulation, and RACK1 is not required for ribosomal localization to growth cones. (A) RACK1 colocalizes with ribosomes, but this colocalization does not increase following BDNF stimulation. In composite images, ribosomes are shown in green and RACK1 is shown in red. Arrowheads in composite images designate regions of colocalization. Dotted lines outline axonal growth cones and include the axon, but axons were not included in the analyses. Note that the brightness and contrast were evenly adjusted throughout the entirety of each image, so that puncta would be easily visible. Scale bars, 5 µm. (B) Quantification of colocalization between RACK1 and ribosomes using Pearson’s correlation coefficient (PCC). *p* = 0.97, Student’s t-test. Vehicle, *n* = 52; BDNF, *n* = 53. (C) Neurons expressing either non-silencing control shRNA or RACK1 shRNA were labeled for ribosomal protein s6 using ICC. Dotted lines outline axonal growth cones. Note that dotted lines also include the axon for ease of visualization, but axons were not included in the analyses. Scale bars, 5 µm. (D) Quantification of ribosomal localization in growth cones. The fluorescent signal of growth cones expressing RACK1 shRNA were normalized to non-silencing control shRNA expressing growth cones. *p* = 0.09, Mann-Whitney. Non-silencing control shRNA, *n* = 79; RACK1 shRNA, *n* = 76.

Because RACK1 is a ribosomal scaffolding protein, we hypothesized that RACK1 knockdown may decrease ribosomal localization to growth cones. Cortical neurons were transfected with either non-silencing control shRNA or RACK1 shRNA. Cells were fixed, and quantitative immunocytochemistry was performed to label ribosomal protein s6, a component of the 40S ribosomal subunit. Measurements of fluorescence intensity within growth cones of pyramidal neurons demonstrated that ribosomal localization to growth cones is unaffected by RACK1 knockdown (Figure 3, C-D). Thus, although RACK1 binds to ribosomes in growth cones, it is not required for ribosomal localization to growth cones.

Although we find that RACK1 is not required for ribosomal localization to growth cones, it is likely that its ribosomal binding function is required for various aspects of neural development. Previously, we showed that RACK1 expression and phosphorylation regulate point contact formation in growth cones (8). Here, we asked specifically whether the ribosomal binding function of RACK1 regulates point contact formation in growth cones under basal and BDNF-stimulated conditions. To investigate this possibility, cortical neurons were transfected with either a His-Myc-tagged empty construct (Myc-control), a His-Myc tagged RACK1 wild type construct (Myc-RACK1-WT), or a His-Myc tagged RACK1 construct that is unable to bind to ribosomes (Myc-RACK1-DE) (29). Cells were fixed, and quantitative immunocytochemistry was performed to label paxillin, a marker of point contacts. Consistent with our previous experiments (8), BDNF stimulation significantly increased point contact density within growth cones. However, overexpression of Myc-RACK1-WT or Myc-RACK1-DE eliminates this BDNF-induced increase (Figure 4, A-B). We had previously shown that appropriate levels of RACK1 are critical for point contact density (8) and here we show that translation mediated by RACK1 likely also regulates BDNF-induced point contact formation in growth cones.

**Figure 4:**
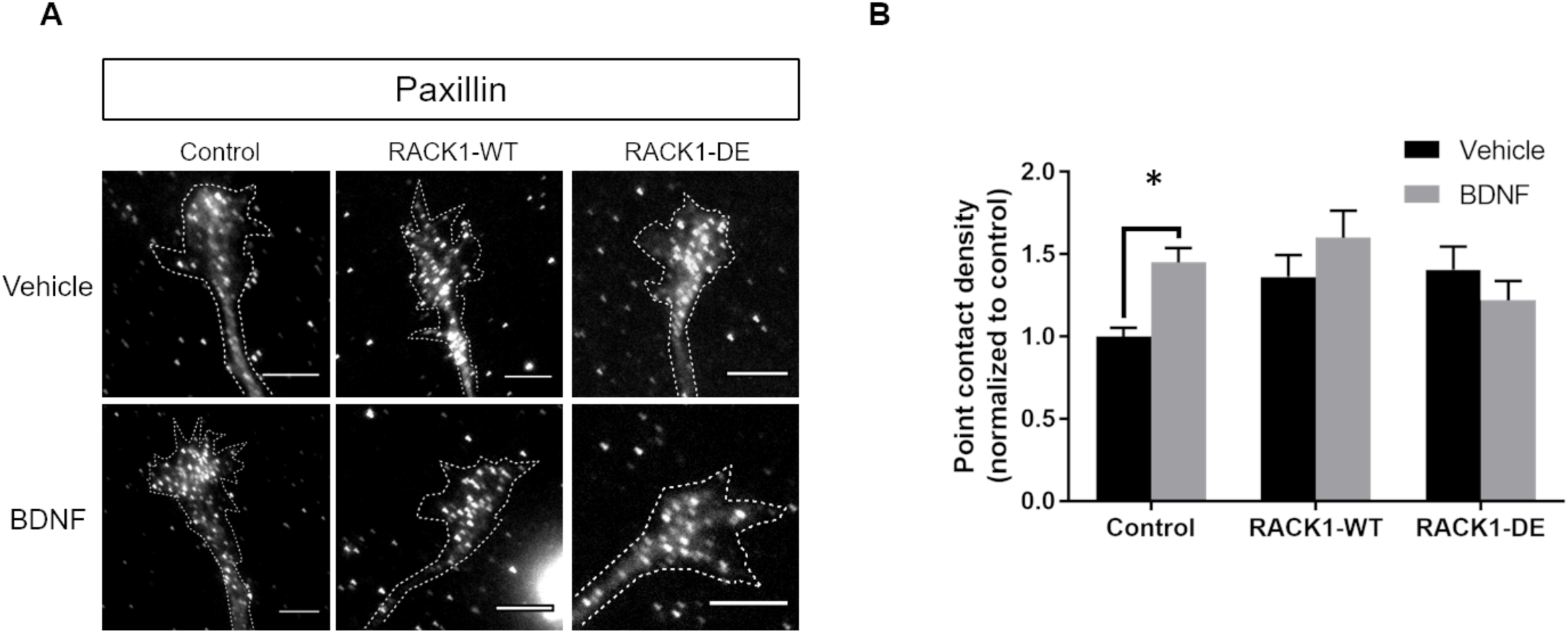
The ribosomal binding function of RACK1 regulates point contact density. (A) Immunocytochemistry for paxillin demonstrates that a BDNF-induced increase in point contact density requires appropriate levels of RACK1 and the ribosomal binding function of RACK1. Dotted lines outline axonal growth cones and include the axon, but axons were not included in the analyses. Note that the brightness and contrast were evenly adjusted throughout the entirety of each image, so that puncta would be easily visible. Scale bars, 5 µm. (B) There is a significant increase in point contact density (puncta/µm^2^) following BDNF stimulation in control neurons. Overexpression of Myc-RACK1-WT or RACK1-DE eliminates the increase in point contact density following BDNF stimulation. p ≤ 0.05, Two-way ANOVA with Dunnett’s multiple comparisons. Myc-control + vehicle, *n* = 50; Myc-control + BDNF, *n* = 55; Myc-RACK1-WT + vehicle, *n* = 56; Myc-RACK1-WT + BDNF, *n* = 52; Myc-RACK1-DE + vehicle, *n* = 52; Myc-RACK1-DE + BDNF, *n* = 52.

Given that the ribosomal binding function of RACK1 is required for BDNF-induced point contact formation, we next asked whether the ribosomal binding function of RACK1 is required for functional aspects of neuronal development, such as axon outgrowth and guidance. We have previously shown that RACK1 is required for axon outgrowth (8), and here we show that RACK1 is required for axon guidance (Figure 1). We expect the underlying mechanism is the ribosomal binding function of RACK1 because RACK1 is required for BDNF-induced translation of β-actin in growth cones (Figure 2). To assess effects on axon growth, cortical neurons were transfected with either Myc-control, Myc-RACK1-WT, or Myc-RACK1-DE, as in previous experiments, and axon length was compared between groups. Overexpression of Myc-RACK1-WT or Myc-RACK1-DE significantly impairs axon outgrowth versus Myc-control (Figure 5, A-B). Thus, RACK1 is critical for axon outgrowth.

**Figure 5:**
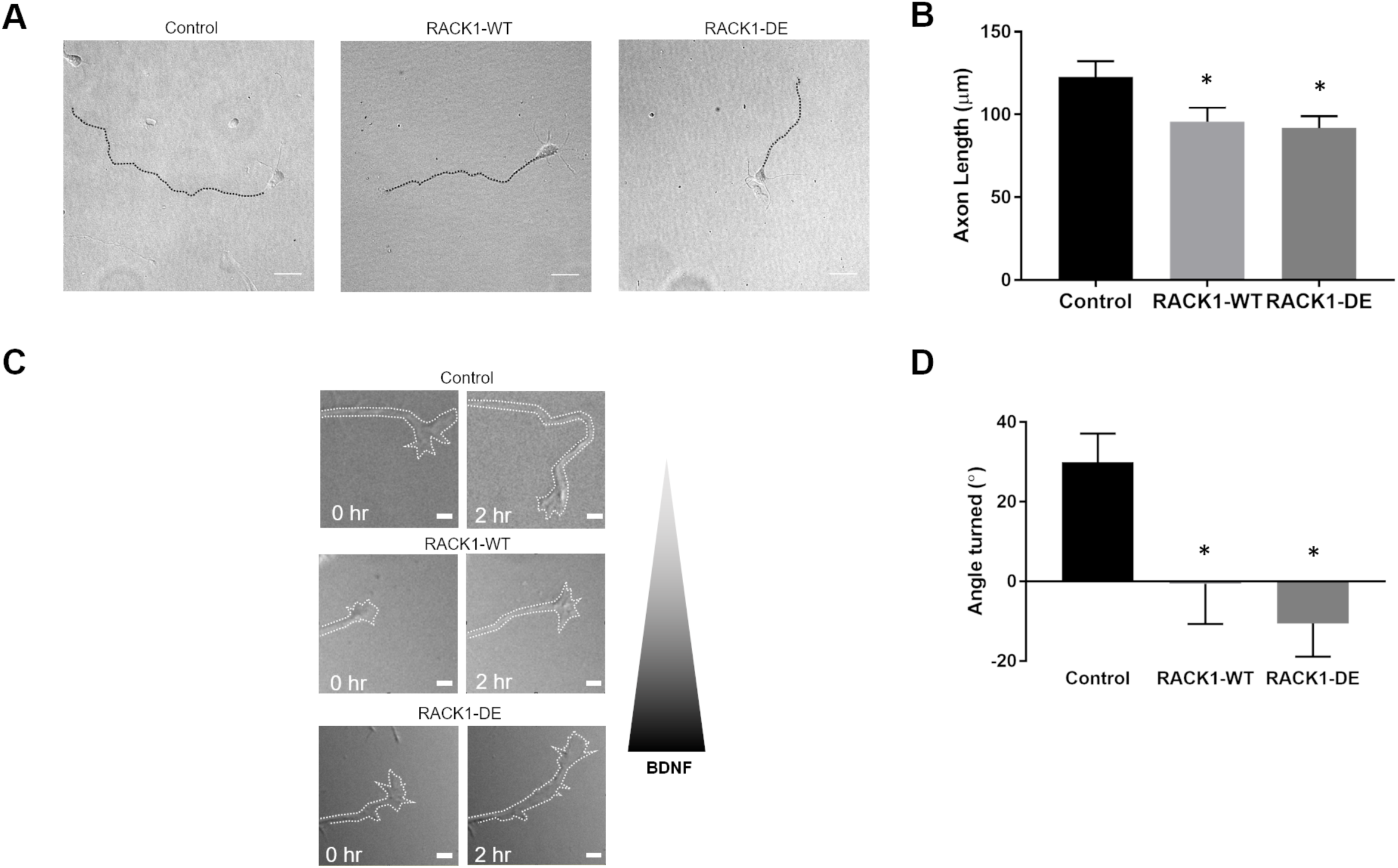
The ribosomal binding function of RACK1 regulates axon growth and guidance. (A and B) Neurons expressing either Myc-control, Myc-RACK1-WT, or Myc-RACK1-DE were fixed at 3 DIV, and axon length was measured. Dotted lines denote the axon in each DIC image. Scale bars, 25 µm. \**p* ≤ 0.05, ANOVA with Dunnett’s multiple comparisons. Myc-control, *n* = 57; Myc-RACK1-WT, *n* = 58; Myc-RACK1-DE, *n* = 57. (C and D) Cortical neurons expressing either Myc-control, Myc-RACK1-WT, or Myc-RACK1-DE were used in the Dunn chamber turning assay. The outer well of the Dunn chamber contained BDNF, which created a chemoattractive gradient. The triangle to the right of the images indicate the BDNF gradient, such that highest concentrations of BDNF occur at the bottom of each image. Turning angles towards the gradient were measured for the three groups. Scale bars, 5 µm. \**p* ≤ 0.05, Student’s t-test. Myc-control, *n* = 14; Myc-RACK1-WT, *n* = 12; Myc-RACK1-DE, *n* = 10.

To investigate whether the ribosomal binding function of RACK1 is required for axon guidance, cortical neurons were transfected with either Myc-control, Myc-RACK1-WT, or Myc-RACK1-DE, and the Dunn chamber axon guidance assay was performed, as in Figure 1. We find that the ribosomal binding function of RACK1 regulates axon guidance. Overexpression of Myc-RACK1-DE eliminates growth cone turning in response to a BDNF gradient (Figure 5, C-D). Taken together, our experiments show that local translation mediated by RACK1 likely regulates axon growth and guidance. Furthermore, they strongly suggest appropriate neural development depends on the regulation of local translation by RACK1. Taken in context of our previous data showing that RACK1 is necessary for the BDNF-induced formation of point contacts (8), this also suggests that local translation at point contacts may be the underlying mechanism.

### Local translation occurs at point contacts

Local translation of certain mRNAs, and specifically β-actin mRNA, in growth cones is necessary for axon guidance (9, 30). We recently found that RACK1 and β-actin mRNA are localized to point contacts and this localization increases following BDNF stimulation (8). Point contacts regulate axon growth and guidance (15, 16, 18, 19). Taken together, these data suggest that local translation occurs at point contacts. Here, we sought to directly demonstrate that point contacts are sites of local translation within growth cones.

We first examined the colocalization between β-actin protein and point contacts in mouse cortical neurons, following either vehicle or BDNF stimulation. Double label quantitative immunocytochemistry to label β-actin protein and paxillin was performed. We then determined the Pearson’s correlation coefficient between β-actin protein and point contacts within growth cones of cortical pyramidal neurons. We find that β-actin protein and point contacts colocalize, but this colocalization does not increase following BDNF stimulation (Figure 6, A-B). We conclude that this is likely due to the abundance of β-actin protein in growth cones, which is certainly not entirely nascent protein resulting from the 30-minute BDNF stimulation. Thus, the small population of newly translated β-actin protein resulting from BDNF is likely overwhelmed by the existing β-actin protein, and it is expected that both populations colocalize with point contacts.

**Figure 6:**
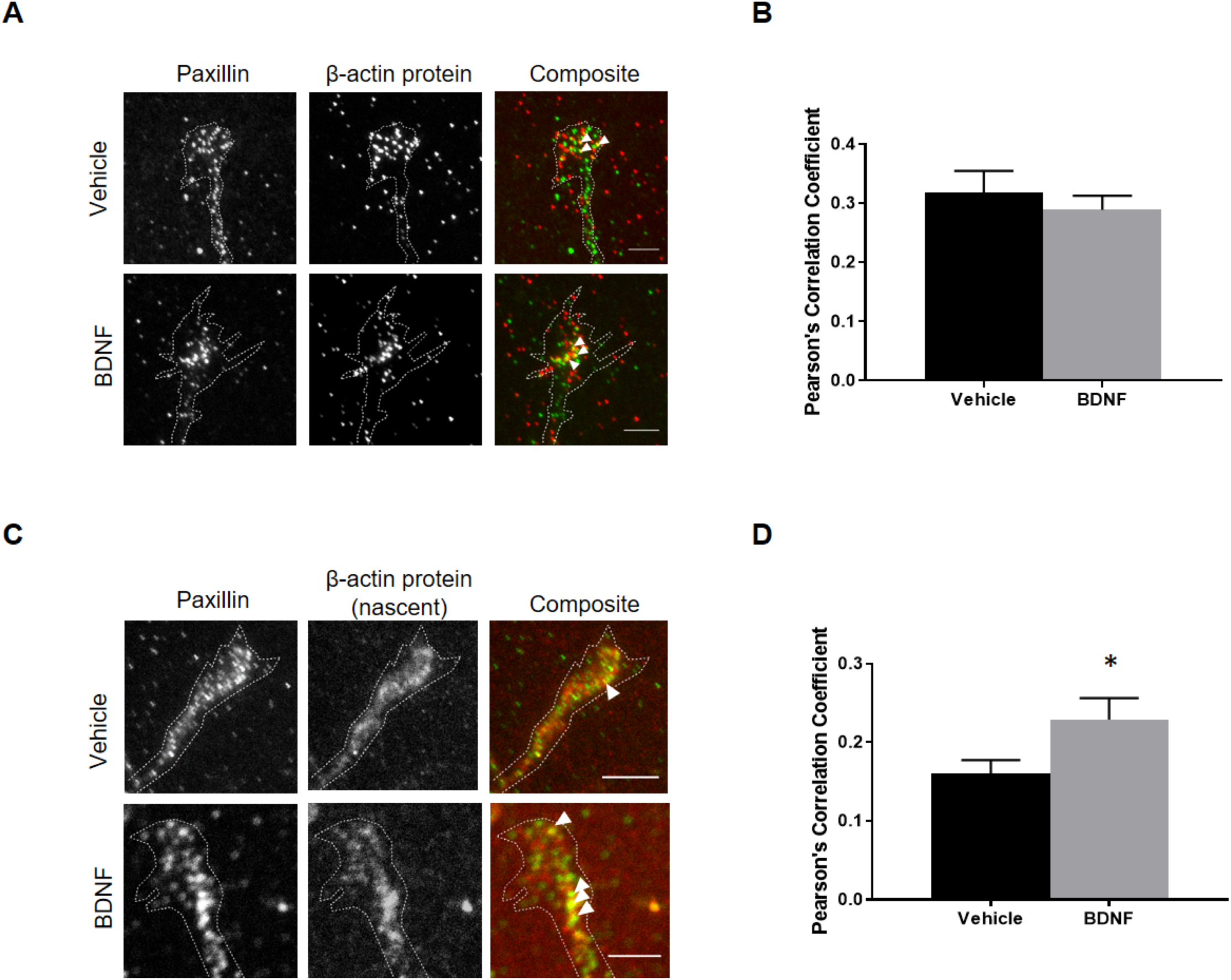
β-actin is locally translated at point contacts. (A) β-actin protein colocalizes with paxillin, a marker of point contacts, but this colocalization does not increase following BDNF stimulation. In composite images, paxillin is shown in green and β-actin protein is shown in red. Arrowheads in composite images designate regions of colocalization. Dotted lines outline an axonal growth cone. Note that dotted lines also include the axon for ease of visualization, but axons were not included in the analyses. Scale bars, 5 µm. (B) Colocalization between paxillin and β-actin protein was quantified using Pearson’s correlation coefficient (PCC). *p* = 0.50, Student’s t-test. Vehicle, *n* = 34; BDNF, *n* = 33. (C and D) Neurons expressing Halo-actin were sequentially labeled with JF549 prior to BDNF/Vehicle treatment and JF646 ligands during BDNF/Vehicle treatment to label preexisting and new translation, respectively. Neurons were then fixed, and ICC was performed to label paxillin. Nascent β-actin protein colocalizes with paxillin, a marker of point contacts, and this colocalization increases following BDNF stimulation. In composite images, paxillin is shown in green and β-actin is shown in red. Arrowheads in composite images designate regions of colocalization. Dotted lines outline an axonal growth cone. Note that dotted lines also include the axon for ease of visualization, but axons were not included in the analyses. Scale bars, 5 µm. (D) Colocalization between paxillin and nascent β-actin protein was quantified using Pearson’s correlation coefficient (PCC). \**p* ≤ 0.05, Student’s t-test. Vehicle, *n* = 29; BDNF, *n* = 28.

Because of the limitations of this analysis, we employed a more advanced method to examine localization of only nascent β-actin protein. To accomplish this, we transfected mouse cortical neurons with the Halo-actin translation reporter used in Figure 2. At 2 DIV, cells were incubated with HaloTag ligands to label preexisting and new translation as described previously (Figure 2C). Cells were immediately fixed, and immunocytochemistry was performed to label paxillin. We then determined the Pearson’s correlation coefficient between nascent β-actin protein and paxillin. Interestingly, we show that the localization of nascent β-actin protein at point contacts increases following BDNF stimulation (Figure 6, C-D). In conclusion, these data provide strong evidence that point contacts are sites of local translation in growth cones.

## Discussion

### Regulation of axon guidance and local translation by RACK1

Studies in *C. elegans* demonstrate that RACK-1, the paralog of RACK1, is necessary for axon guidance in invertebrates (6), but the role of RACK1 in mammalian neural development is not well understood. RACK1 is required for neural tube closure (5) and initial neurite formation (7), and RACK1 knockout mice are gastrulation lethal (4). Thus, RACK1 is critical for global mammalian development, but also contributes specifically to neural development. We recently showed that RACK1 is required for functional aspects of nervous system development in mammals, such as axon outgrowth and neurotrophin-induced growth cone spreading (8). These data led us to hypothesize that RACK1 would be required for axon guidance. Here, we show that RACK1 is required for axon guidance in mammals (Figure 1).

It is possible that RACK1 regulates growth cone guidance via its role as a member of the local translation complex because local translation of β-actin mRNA is necessary for appropriate axon guidance (11, 12). RACK1 is a highly conserved ribosomal scaffolding protein and its interaction with ribosomes in non-neuronal cells is well characterized (25). Furthermore, RACK1 binds the β-actin mRNA/ZBP1 complex on ribosomes, and after stimulation, β-actin mRNA is released and translated in a Src-dependent manner (10). Blocking RACK1 phosphorylation by Src via overexpression of a RACK1 mutant, RACK1-Y246F, reduces the translation of β-actin mRNA (10) and impairs axon outgrowth and BDNF-induced growth cone spreading (8). Here, we directly demonstrate that loss of RACK1 reduces β-actin local translation (Figure 2). Though, it is important to note that overexpression of RACK1-Y246F induces accumulation of β-actin mRNA within growth cones because the release of β-actin mRNA is impaired (10). Therefore, if a compensatory mechanism existed to facilitate local translation of β-actin mRNA in the absence of RACK1, it is possible that this would be obscured in the RACK1-Y246 experiments because the mRNA would be sequestered. RACK1 knockdown does not impair local translation of β-actin mRNA in a manner similar to the non-phosphorylatable RACK1 mutant because RACK1 knockdown would not result in the sequestration of β-actin mRNA. Thus, these data suggest that a compensatory mechanism does not exist to facilitate local translation of β-actin mRNA in the absence of RACK1.

Here, we show in both fixed and living cells that RACK1 is required for local translation of β-actin in growth cones (Figure 2). We also find that RACK1 knockdown reduces nascent β-actin protein to levels similar to a control group treated with cycloheximide, a known protein synthesis inhibitor (Figure 2). Taken together, these data suggest RACK1 regulates local translation and is required for axon guidance and by extension, appropriate neural development.

### Ribosomal binding function of RACK1 regulates neural development

In the current study, we show that RACK1 is necessary for axon guidance (Figure 1), and in our earlier study we showed that RACK1 is necessary for axon outgrowth and BDNF-induced growth cone spreading (8). Because RACK1 interacts with many different signaling molecules, including PKC, Src, ribosomes, and integrins (2), disruption in one or more of these pathways may be the cause of the observed deficits in neural development following RACK1 knockdown. Therefore, it is of interest to elucidate which of these pathways is involved in these deficits. Previously, we showed that the interaction between RACK1 and Src regulates axon outgrowth and BDNF-induced growth cone spreading (8). Because Src phosphorylation of RACK1 is required for the local translation of β-actin mRNA (10), we hypothesized that binding between RACK1 and ribosomes is also necessary for neural development as both Src and ribosomes modulate local translation. In addition, our aforementioned experiments demonstrating that RACK1 is required for local translation suggest that this is a plausible mechanism because local translation is required for axon guidance (11, 12).

We first showed that the association between RACK1 and ribosomes does not increase following BDNF stimulation (Figure 3). However, this is not entirely surprising as proteomes of *T. lanuginosus* and *S. cerevisiae* show that most ribosomes are RACK1 bound (31). As RACK1 is a highly conserved protein (25), it is possible that this association between RACK1 and ribosomes extends to all eukaryotes, including mammals. But it is important to note that studies have shown a non-ribosome-bound form of RACK1 in human cells (32), and ribogenesis and ribosomal stability does not require binding between RACK1 and ribosomes (33).

Here, we show that the ribosomal binding function of RACK1 regulates axon outgrowth (Fig. 5A-B) and axon guidance (Fig. 5C-D), but not ribosomal transport to growth cones (Figure 3). Though the importance of RACK1 for neural development has been shown in *C. elegans* (6), the specific RACK1 binding partners that contribute to this regulation is not well understood. Because RACK1 is involved in such a wide variety of signaling cascades, knowing which binding partner modulates neural development has very important implications for both basic research and for potential therapeutic treatments. Interestingly, we show that RACK1 is not required for ribosomal localization to growth cones (Figure 3), suggesting RACK1 does not transport ribosomes to growth cones, or there are compensatory mechanisms for when RACK1 is lost. Another possibility is that ribosomes are themselves locally translated in growth cones, which is not unexpected as mRNAs encoding ribosomal subunits have been observed in growth cones (34). Thus, this suggests that RACK1 does not regulate neural development through ribosomal transport. Here, we show that the ribosomal binding function of RACK1 regulates the BDNF-induced increase in point contact density. It is likely that RACK1 mediated local translation of β-actin acts to stabilize point contacts in the same manner that β-actin stabilizes focal adhesions (35). Thus, it is expected that an absence of RACK1’s translational function, which occurs in overexpression of the RACK1-DE mutant, results in the destabilization of point contacts so that they cannot form, or their lifetime is so shortened that there is no net increase in point contacts following stimulation. Because RACK1 regulates the translation of other proteins besides β-actin, it is also possible that RACK1 is responsible for the local translation of other proteins involved in point contact formation. Previous studies in mouse embryonic fibroblasts and squamous cell carcinoma cells show that RACK1 localizes to nascent focal adhesions (36) and we show RACK1 localizes to point contacts in neurons (37). In the context of these experiments, our new data demonstrate that RACK1 binding of ribosomes regulates the formation of point contacts and functional aspects of neural development such as axon outgrowth and guidance.

### Point contacts are sites of local translation in growth cones

Although it has been well-established that local translation occurs in axons and growth cones, we do not know where this translation occurs within these neuronal compartments. One recent study demonstrated that late endosomes are a platform for local translation within axons (38). Here we add to our knowledge about platforms for local translation by demonstrating that local translation also occurs at point contacts. Point contacts are found exclusively in neuronal growth cones and are somewhat analogous to focal adhesions in other cell types. Given their similarities, studies on focal adhesions in nonneuronal cells have some relevance for the current studies on point contacts in neurons. Although it has not been directly shown, previous studies strongly suggest that focal adhesions are sites of local translation. In mammalian epithelial cells and fibroblasts, integrin binding results in the localization of mRNA and ribosomes to focal adhesions (39). In spreading and migrating human fetal lung fibroblast cells, ribosomes and initiation factors localize at focal adhesions (40). Further, focal adhesion stability and cell migration increases following β-actin mRNA compartmentalization to focal adhesions in mouse embryonic fibroblasts (35). Finally, recent advances in single molecule tracking enabled researchers to show that β-actin mRNA at focal adhesions in mouse embryonic fibroblasts exhibits movement characteristic of increased translation (23). In the current study, we directly show local translation at adhesions by combining translation reporters and point contact markers.

Point contacts are required for axon growth and guidance. They not only dynamically respond to guidance cues (18), but also provide the mechanical force necessary for axon extension (19). Thus, point contacts would be an optimal site for local translation in order to have a maximal impact on axon guidance. Previously, we showed that members of the local translation complex, β-actin mRNA and RACK1, localize at point contacts and that this localization increases following BDNF stimulation (8). These data, in addition to our data showing that the ribosomal binding function of RACK1 underlies neural development and point contact formation, highly suggest that local translation occurs at point contacts and prompted us to further investigate this possibility.

Here, we are the first to demonstrate that point contacts are sites of local translation in growth cones. Given our previous findings that BDNF stimulation increases the localization of β-actin mRNA at point contacts (8), we hypothesized that BDNF stimulation would increase local translation of β-actin mRNA at point contacts. Indeed, we found that BDNF stimulation increases the localization of nascent β-actin protein at point contacts (Figure 6C-D). We note that our initial experiment did not corroborate this finding (Figure 6A-B). However, this is most certainly due to the fact that all β-actin protein in growth cones was labeled rather than nascent β-actin protein alone. Therefore, it is likely that abundant preexisting β-actin confounded the analysis.

The finding that local translation occurs at growth cone point contacts has very important implications for a number of reasons. First, it suggests that local translation and cell adhesion act coordinately to direct neural development. Thus, deficits in local translation may impair cell adhesion and vice versa. Further, it suggests that cell adhesion molecules as a whole may be critical for local translation. Therefore, in future experiments we will examine the effect of knocking down cell adhesion molecules on local translation. In line with this idea, one study has demonstrated that the efficiency of local translation can be altered by the isoform of laminin that is present (41). In addition, because local translation is not limited to neurons, these findings could have important implications for multiple cell types. Finally, this finding warrants further study into how aberrant cell adhesion molecule expression might contribute to specific developmental disorders via their control of local translation. It is hoped that further understanding the link between cell adhesion and local translation will aid in the development of treatments for developmental and neurological disorders.

## Materials & Methods

### Cell Culture and Transfection

All experimental procedures were approved by the Institutional Animal Care and Use Committee at Kent State University. C57BL/6J mice (The Jackson Laboratory) were bred to produce timed pregnant animals. The day of the plug was considered embryonic day 0 (E0). Cortices were dissected from E17 embryos of either sex. Dissociated cortical cells were plated in MEM with 10% FBS on acid-rinsed coverslips (Carolina Biological) or live cell dishes (MatTek) pretreated with 100 µg/mL poly-L-lysine (Sigma-Aldrich) and 10 µg/mL laminin (Life Technologies). After cells adhered to the coverslips (approximately 2 h), the media was changed to Neurobasal with Glutamax and B27 (all Life Technologies) and cells were cultured for 2 DIV at 37°C in 5% CO2. For transfections, neurons were transfected prior to plating using the Lonza Amaxa™ 4D-Nucleofector™ and P3 Primary Cell 4D-Nucleofector® X Kit with program code CU-133. The nonsilencing control shRNA and RACK1 shRNA were purchased from GE Dharmacon and used at a concentration of 1 µg DNA per transfection or 0.4 µg DNA per transfection if used in combination with other constructs (see the “shRNA Constructs” section for additional details). The nonsilencing control siRNA and RACK1 siRNA were purchased from Santa Cruz and GE Dharmacon respectively and used at a final concentration of 300 nM. The sequence of the RACK1 siRNA is the most optimal computationally generated sequence as reported by the Dharmacon siDESIGN center, and it is identical to the RACK1 shRNA sequence with the exception of 3’ overhanging UU dinucleotides for maximal siRNA effectiveness. The following constructs were used in these experiments: Myc-RACK1-WT, Myc-RACK1-DE (both provided by Dr. Marcello Ceci (29) and both used at a concentration of 0.5 µg DNA per transfection), a control empty Myc vector subcloned from the Myc-RACK1-WT construct (0.5 µg DNA per transfection), and HaloTag β-actin (provided by Dr. Robert H. Singer (27), 0.4 µg DNA per transfection). When comparing basal and stimulated conditions, cells were starved by removing B27 from the medium for 3 h and then treating with 100 ng/mL BDNF (PeproTech) or vehicle control for 30 min at 37°C. In all fixed cell experiments, cells were fixed with 4% paraformaldehyde in phosphate buffered saline. When performing ICC for β-actin protein, a fixation step using 50% ice cold methanol/PBS was added following paraformaldehyde fixation.

### shRNA Constructs

Commercially available shRNA for RACK1 and a nontargeting control shRNA were purchased from GE Dharmacon’s GIPZ construct library. The human cytomegalovirus promoter (hCMV) drives the expression of the shRNA. The GIPZ constructs also contain GFP as a marker of shRNA expression. Only cells clearly expressing GFP were included in analyses. The shRNA was previously validated in neurons (8). The mature antisense sequence used to knockdown RACK1 was: (V3LMM_447148): TCTGAATGACTCTCATCCT. The non-targeting mature antisense sequence was: CTTACTCTCGCCCAAGCGAGAG.

### Immunofluorescence Experiments (IF)

IF experiments were performed as previously described (42), and the following primary antibodies were used: mouse anti-paxillin antibody (1:500, BD Biosciences), rabbit anti-RACK1 antibody (1:500, Santa Cruz Biotechnology), mouse anti-RACK1 antibody (1:500, Santa Cruz Biotechnology), rabbit anti-ribosomal protein S6 antibody (1:100, Santa Cruz Biotechnology), mouse anti-ribosomal protein S6 antibody (1:100, Santa Cruz Biotechnology), mouse anti-β-actin (1:1500, Abcam), and rabbit anti-Myc (1:500, Abcam). The following secondary antibodies were used: goat anti-rabbit Alexa 488, goat anti-mouse Cy3, donkey anti-mouse Alexa 647 (all from Jackson ImmunoResearch), goat anti-mouse Alexa 488, goat anti-rabbit Alexa 488, and goat anti-mouse Alexa 555 (all from Thermo Fisher).

### Image acquisition and analysis

All fixed cell experiments were conducted in a blinded fashion by assigning random numbers as the labels for both the slides and the acquired images. Experiments were only unblinded following analysis. Immunofluorescence was visualized on a Nikon Eclipse Ti2 microscope using either a Nikon S Plan Fluor ELWD 20X (0.45 NA) or a Nikon Plan Apo Lambda 100X (1.45 NA) objective. Images were captured with a Hamamatsu ORCA-Flash4.0 V3 Digital CMOS camera using Nikon NIS elements software. To quantify fluorescence, all experimental and image acquisition parameters were kept constant throughout each experiment. Any images containing overexposed pixels were excluded from analysis.

Axon length, growth cone area, point contact density, and fluorescence intensity were quantified using ImageJ tools within Fiji software (43). We can reliably define the axon as the longest neurite, which extends at least three times the length of the next longest neurite. Axon length was determined by measuring the length of the primary axon (i.e., longest neurite) from the cell body to the center of the axonal growth cone. Axon length does not include the length of any branches coming off the main axon. Growth cones were defined based on images taken in differential interference contrast (DIC). The brightness and contrast were adjusted in each image following acquisition to easily visualize the entire growth cone, including the central domain, lamellipodia and filopodia. Growth cone area was then calculated as the area defined by enclosing the entire perimeter of the growth cone including central domain, lamellipodia, and filopodia, but not the axon. This method is in line with previous studies (18, 42). Point contact density was determined by counting the number of fluorescently labeled paxillin puncta within a growth cone and then dividing that value by growth cone area. Point contacts were counted by hand in growth cones with few puncta; otherwise, they were counted using the particle analysis tools in Fiji (43). A preliminary analysis was performed to verify that the puncta counts from the computerized particle analysis method did not differ from puncta counted by hand. To determine the mean fluorescent intensity in a ROI, the structure of interest (e.g., growth cone) was first outlined. The mean fluorescent intensity was background subtracted in each image by measuring the fluorescent intensity in a region near the ROI but excluding the ROI. Fiji determines mean fluorescent intensity by measuring the total fluorescent intensity and dividing by the area to normalize for differences in ROI size. Pearson’s correlation coefficients (PCC) of a region of interest (ROI) were quantified using the Coloc2 plugin of Fiji software (43). Only cells with a pyramidal morphology were included in these analyses. Each experiment was repeated a minimum of three times.

### Axon guidance assay using Dunn chamber

For Dunn chamber assays, cortical neurons were grown on 22×22 mm square coverslips (Assistent). At 2 DIV, these coverslips containing transfected cells (nonsilencing control shRNA, RACK1 shRNA, empty myc vector, Myc-RACK1-WT, or Myc-RACK1-DE) were placed onto Dunn chambers (Hawksley) and a BDNF gradient was established by adding Hibernate E low fluorescence media (Brain Bits) and a final concentration of 100 ng/mL BDNF (PeproTech) to the outer well of the Dunn chamber; Hibernate E low fluorescence media and vehicle (0.1% BSA) was added to the inner well of the Dunn chamber. Dental wax (VWR) was used to seal the chamber. Images were taken every 5 minutes for 2 hours. Protocols for chamber setup, imaging, and data analysis were conducted as reported previously (24, 44). Imaging conditions of 37°C and 5% CO_2_ were maintained using a Tokai Hit™ incubation system.

### HaloTag β-actin JF549-JF546 Labeling Assay

At 2 DIV, live cell dishes containing cells transfected with HaloTag β-actin and either nonsilencing control siRNA or RACK1 siRNA were starved for 1.5 hours in Neurobasal with Glutamax. Afterwards, JF549 HaloTag ligand (Dr. Luke Lavis, HHMI Janelia Research Campus) was added at a 20nM final concentration in Neurobasal with Glutamax for one hour. Then, five quick washes with HBSS were performed and cells were incubated for 30 minutes in HBSS. Next, JF646 HaloTag ligand (Dr. Luke Lavis, HHMI Janelia Research Campus) was added at a 200nM final concentration in Neurobasal with Glutamax, B27, and 100 ng/mL BDNF for 30 minutes. Finally, five quick washes with HBSS were performed and cells were incubated for 30 minutes in HBSS with 100 ng/mL BDNF. For the nonsilencing control siRNA and cycloheximide (CHX) treatment group, CHX (Calbiochem) was added at a 60 µM concentration alongside BDNF treatment and included in all subsequent steps until the last wash step before imaging. Media was changed to Hibernate E low fluorescence (Brain Bits) for imaging. Imaging conditions of 37°C and 5% CO_2_ were maintained using a Tokai Hit™ incubation system. Only cells clearly showing JF549 staining were included in the analyses.

### Statistical Analysis

All statistical analyses were conducted in GraphPad Prism. Significance was set at *P* ≤ 0.05. The specific statistical test used for each experiment is given in its figure legend. Error bars represent the standard error of the mean from a minimum of three independent experiments unless otherwise indicated.

## Abbreviations

BDNF: Brain-derived neurotrophic factor
IF: Immunofluorescence
RACK1: Receptor for activated C kinase 1
RBP: RNA-binding proteins
UTR: Untranslated region
ZBP1: zipcode binding protein 1

## Acknowledgements

We thank Dr. Marcello Ceci (Tuscia University) for the RACK1 constructs, Drs. Robert Singer and Young J. Yoon (Albert Einstein College of Medicine) for the HaloTag β-actin construct, and Dr. Luke Lavis (HHMI Janelia Research Campus) for the HaloTag ligands.

